# The Effect of Blood Viscosity on Shear-Induced Hemolysis using a Magnetically Levitated Shearing Device

**DOI:** 10.1101/2021.08.02.454383

**Authors:** James A. Krisher, Richard A. Malinauskas, Steven W. Day

## Abstract

**Introduction:** Blood contacting medical devices, including rotary blood pumps, can cause shear-induced blood damage that may lead to adverse effects in patients. Due in part to an inadequate understanding of how cell-scale fluid mechanics impact red blood cell membrane deformation and damage, there is currently not a uniformly accepted engineering model for predicting blood damage caused by complex flow fields within ventricular assist devices (VADs).

**Methods:** We empirically investigated hemolysis in an axial Couette flow device typical of a rotary VAD to expand our current understanding of shear-induced blood damage in two ways. First, we used a magnetically levitated device to accurately control the shear rate and exposure time experienced by blood and to minimize the effects of other uncharacterized stresses. Second, we explored the effects of both hematocrit and plasma viscosity on shear-induced hemolysis to characterize blood damage based on the viscosity-independent shear rate, rather than on shear stress.

**Results:** Over a shear rate range of 20,000-80,000 1/s, the Index of Hemolysis was found to be largely independent of hematocrit, bulk viscosity, or the suspension media viscosity.

**Conclusion:** It is recommended that future investigations of shear-induced blood damage report findings with respect to the viscosity-neutral term of shear rate, in addition to the bulk whole blood viscosity measured at an appropriate shear rate relevant to the flow conditions of the device.

## Introduction

Current ventricular assist devices (VADs), as well as other blood contacting medical devices, can exert magnitudes of fluid shear that exceed physiological levels. This can ultimately result in damage to blood components including red blood cells (RBCs), platelets, and von Willebrand factor, and result in adverse patient events such as bleeding, thrombosis, and renal injury. Quantitative relationships between applied fluid shear, exposure duration, and resulting cell damage have been developed based on empirical fits to data based on specific flow devices and test conditions. However, the mechanistic understanding of shear-induced damage that considers cell biomechanics and cell-scale fluid mechanics is inadequate to generate quantitative thresholds that are predictive of damage caused by complex flow fields within VADs. We investigated hemolysis in an axial-flow model representative of some rotary VADs to expand our current understanding of shear-induced blood damage in two ways. First, we used a novel magnetically levitated rotor device to accurately control shear stress experienced by blood without introducing uncharacterized stresses, such as the significant stresses from conventional friction bearings. Second, we explored the effects of both plasma viscosity and hematocrit on shear-induced hemolysis to evaluate the ramifications of characterizing blood damage based on shear stress, a parameter which depends on blood viscosity, versus viscosity-independent shear rate.

### Hemolysis as a measure of shear-induced damage

One of the most commonly used surrogates for shear-induced blood damage is hemolysis, as typically quantified by an increase in the concentration of plasma free hemoglobin. An RBC can be mechanically deformed by fluid shear to the point of stretching pores in its membrane, or to a lethal rupture, leading to the release of cellular contents [1]. There are several important reasons to elucidate the mechanisms leading to shear-induced hemolysis: 1) Clinical markers for hemolysis (plasma free hemoglobin, serum lactate dehydrogenase, hemoglobinuria, anemia), are commonly monitored as they have been linked to increased adverse patient events and mortality [2]–[4]; 2) The degree of hemolysis caused by a new blood pump is typically compared to that of a comparator clinical device throughout its development and evaluation to assess design changes, functional reproducibility between devices, and safety [5][6]; 3) The measurement of plasma free hemoglobin is an accessible and well-validated procedure with a high degree of resolution and, with the additional measurement of hematocrit and total hemoglobin, can be normalized between blood pools allowing for potential comparisons to be made [7]; And 4), as shear flows can affect other components in blood, including those involved in thrombus formation and coagulation, the extent of RBC damage serves as a surrogate for fluid-induced mechanical effects to other blood elements.

### The type of flow in test devices matters

Numerous experiments have been conducted by different research groups in an effort to quantitatively relate applied fluid shear stress (τ, Pa) and blood cell exposure time (t_exp_, ms) to the resulting hemolysis through a power-law relationship (%Index of Hemolysis = %IH = C τ^α^ t_exp_^β^, where α, β, and C are experimentally determined constants) [8][9][10]. While the power-law equation is commonly referenced, researchers have pointed out that the coefficients used in this equation to predict hemolysis vary among testing laboratories and devices, and when using blood from different species [11]–[13]. Hence, there is not a consensus regarding a predictive fluid shear/ blood damage index equation for use to enhance development and the safety evaluation of devices. As concluded in recent review articles [14][15], new experiments that accurately represent the flow fields encountered in cardiovascular devices are necessary to inform the development of a broadly applicable hemolysis prediction model.

Given the importance of the flow regime and fluid parameters to the study of shear-induced hemolysis, certain relevant terms must be clearly identified. Blood damage studies have employed devices which apply fluid shear via three categories of flow: (i) Couette/Taylor-Couette flow, created in rheometers and some rotary VADs; (ii) Poiseuille flow, common in needles and micro-fluidic blood damage studies; and (iii) turbulent flow, attainable in nozzles and needle models [16] [17]. Wurzinger, et al. [18] and Heuser and Opitz [19] used a Couette viscometer to expose cells to shear stresses for relatively short durations (0.01-1 second), while Leverett [9] investigated longer (>100 seconds) durations. They measured the shear stress required to release 1 percent of the RBC’s hemoglobin as a function of exposure time and reported a noticeably higher sensitivity of the red cells to shearing forces than had been previously shown. More recently, Paul, et al. [20] used a similar Couette viscometer, but employing a fluid seal, and reported lower hemolysis levels (higher threshold stresses) than Wurzinger [21]. Zhang, et al. [22] modified an axial flow VAD by removing its rotor blades to create a shearing device in which the RBC exposure duration could be controlled, but their device contained mechanical bearings. Conventional mechanical bearings (e.g. pin bearings) may cause unaccounted for blood damage that is secondary to the primary rotor shear forces intended to be controlled and interrogated in a given study. This study details the design and utilization of a novel axial-flow magnetically levitated shearing device capable of applying controlled fluid shear to blood for regulated durations. The use of touch-free magnetic bearings minimizes uncharacterized sources of damage such as those of conventional mechanical bearings. Our shearing device applies a Taylor-Couette flow similar to the flow fields found in some modern axial-flow rotary blood pumps.

### Fluid stress and RBC Mechanical Strain

Blood consists of a suspension of cells, primarily RBCs (typically 36-50% by volume), in a protein rich aqueous plasma. Human RBCs measure 6-8 μm in diameter by 2.2 μm in thickness with a distinctive biconcave disc shape. Their inherent deformability allows them to maneuver through small diameter blood vessels by contorting into a number of configurations without straining the cell membrane beyond its damage threshold. An RBC experiences a mechanical strain of its cell membrane that is a function of the fluid stress, including shear, exerted by the surrounding fluid over some length of time. According to a theory first proposed by Rand [23], the RBC membrane becomes compromised (pore formation or tearing) and releases cellular material (i.e. hemolysis) when the mechanical strain of the cell membrane reaches a threshold value. Although the terms stress and strain are sometimes imprecisely used interchangeably, the strain of the membrane is related to the stress within the membrane and is governed by the membrane material properties. Additionally, the relationship between membrane stress and strain is time dependent due to the viscoelastic nature of the cell membrane [23]. As a result, cells can survive exposure to high levels of stress for short time durations better than they can for longer exposure times, hence the development of power-law models of hemolysis considering stress and exposure time [9][24].

What is less clear is the relationship between the stress within the fluid surrounding an RBC and the strain within the cell membrane. Since cell shape and orientation can vary and depend on the fluid flow, strain, and shear, the membrane stress does not need to be equal or even directly proportional to the surrounding fluid shear stress. Moreover, the effect of fluid shear and extensional stresses may affect cell membrane strain very differently [25][14], [24]. As a simple example, a cell that rolls within a shear flow will experience membrane stresses lower than the bulk fluid stress because the velocity gradient experienced by the cell is diminished in the cell’s rotating reference frame. Despite this, most experimental studies and predictive models for hemolysis, including most of those cited here, have used fluid shear stress based on the bulk fluid viscosity as a principle parameter. This assumes that the stress within the cell membrane, which ultimately leads to leakage of the cell contents, is proportional to the bulk fluid stress. This is a flawed mechanistic model, as explained by Faghih, et al. [14], but is an inherent assumption made when using the most pervasive method (i.e., the power-law model) of estimating shear induced blood damage.

### Fluid shear stress is dependent on viscosity and shear rate

The use of fluid shear stress as a principle determinant of blood damage is further complicated because of a range of definitions for the viscosity of a suspension, such as blood. Within a fluid, shear stress, *τ,* is defined as the product of viscosity, *μ,* and the gradient of velocity, known as the fluid strain rate or shear rate, 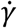 according to 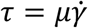. This requires a characterization of both applied shear rate and blood viscosity (μ). However, the viscosity of whole blood is a function of the hematocrit, plasma viscosity, and the shear rate. While plasma has a nearly constant viscosity across a wide range of shear rates, bulk blood viscosity decreases with shear rate as RBCs separate and undergo deformation due to the velocity gradient [26]. The viscosity reaches an asymptotic value as the cells become maximally separated and deformed, thus blood viscosity dependence on shear rate is most pronounced below 1,000 1/s [27]. We refer to three different viscosities within this study to describe the non-Newtonian nature of blood and relate it to bulk and local shear-induced blood damage. We use the term *low-shear bulk viscosity* to characterize the whole blood viscosity at low shear rates (<1,000 1/s) such as those measured in a capillary viscometer. The second is *high-shear bulk viscosity* defined as the bulk blood viscosity in a relevant high shear flow environment, such as in a VAD (shear rate > 10,000 1/s). High-shear bulk viscosity is always lower than low-shear bulk viscosity due to the shear thinning non-Newtonian nature of blood. Lastly, we refer to the *media viscosity* which, for plasma in the case of whole blood, acts as a Newtonian fluid with a nearly constant viscosity under both low and high fluid shear conditions.

To evaluate the effect of fluid shear stress on blood damage, we must consider the factors described above and how they relate to the forces the fluid exerts on a suspended cell and the resulting membrane strain that cell experiences. For laminar flows at high shear rates and typical blood hematocrits, mechanical stress on the cell membrane is the result of cell-media interactions [27], [28]. While mechanical stress may also result from cell-cell interactions, these are thought to be most significant for turbulent flows [29] or at very low shear rates [27]. This study examines the impact of media viscosity and bulk viscosity on shear-induced blood damage in a laminar flow device, and the value of using the viscosity-independent term shear rate for damage characterization rather than the commonly used viscosity-dependent term shear stress. We tested this relationship by varying the bulk viscosity of blood in two ways. First, by adjusting hematocrit, we altered the bulk viscosity while maintaining a near constant media (plasma) viscosity. Second, we created blood samples with similar hematocrits but adjusted the viscosity of the plasma suspension media using Dextran-40, which consequently changed the viscosity of the bulk whole blood. The hematocrit and viscosity adjusted blood were exposed to a range of supraphysiological shear flows and the resulting red blood cell damage evaluated. In this manner, these experiments effectively decouple shear rate from shear stress in order to explore the dependence of cell damage on shear rate and viscosity in a flow regime that is relevant to the design of VADs and other blood contacting devices.

## Materials and Methods

### Device Characterization

This investigation used an axial flow, single pass, Couette shear device to apply a controlled fluid shear to blood for a set duration (Figure 1). A comprehensive description of the device and its characterization may be found in Krisher [30]. The device was constructed by modifying an existing axial flow VAD developed at the Rochester Institute of Technology, the Lev-VAD [31], the novel feature of which is an active magnetic bearing system used to levitate the pump’s titanium impeller [32]. The impeller vanes were ground off and the vaneless rotor body was fitted with a tapered Delrin sleeve located at the center of the rotor. The magnetically-levitated (mag-lev) rotor in the shearing device allowed for blood to be exposed to a controlled fluid shear application in a single “shearing gap” region with no uncharacterized shear contributed by conventional friction bearings. Inserts at the inlet and outlet of the device were used to reduce the fluid volume to 6 ml and to facilitate connection via Luer fittings. A syringe pump was used to provide a controlled flow rate of blood through the device.

**Figure 1.**
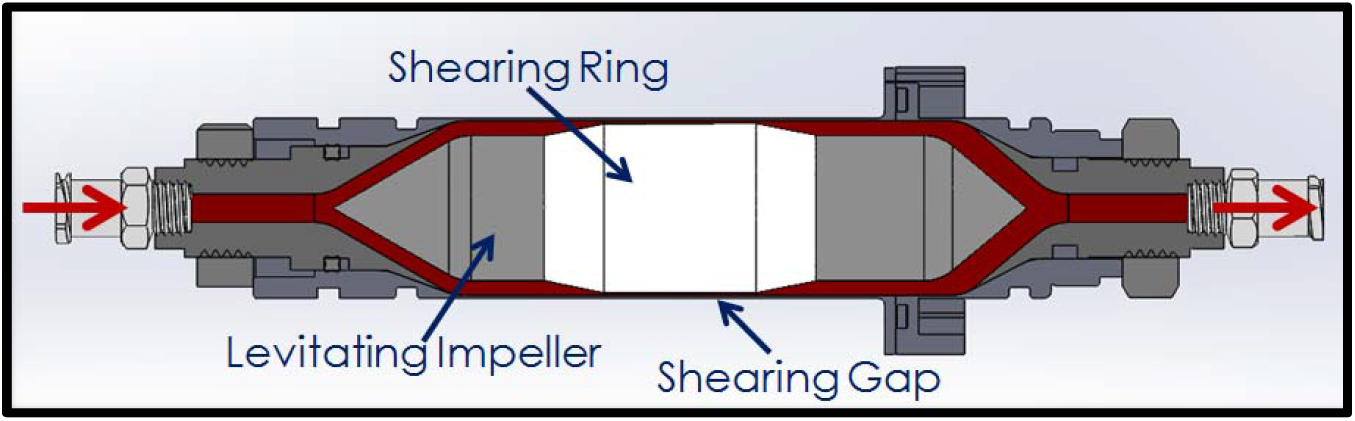
Cross section of shearing device. Entrance to exit length is approximately 15 cm. The grey blood contacting elements are the titanium housing and impeller, while the white Delrin impeller sleeve creates the shearing gap. The blood pathway is shown in red.

The primary parameters of interest in this investigation are the magnitude of applied fluid shear, the exposure time to that shear stress, fluid viscosity, and shear rate. Shear rate (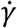, 1/s) is classically defined in Couette flow as the velocity gradient between two surfaces across a fluid gap. In the Taylor-Couette flow for this device, with a rotating inner cylinder and stationary outer cylinder, the shear rate is a function of the radius (R) of the inner cylinder (the shearing ring in our case), the width of the annular fluid gap (a), and the rotor speed in RPM. Exposure time (t_exp_, ms) is calculated as a function of the volumetric flowrate through the device (Q) and the geometry of the shearing region (cross sectional area of the gap = A_CS_).

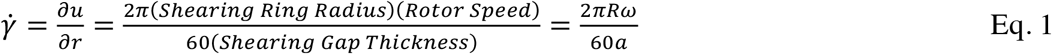

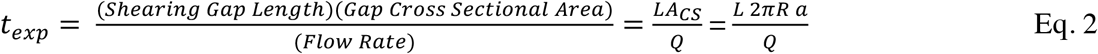

In this study, the annular shearing gap was 127 μm thick and 13 mm long (L); the geometry of the flow path is shared as supplementary material to this article. The rotor maintained stable levitation in blood between 2,500 RPM and 12,000 RPM, which allowed a shear rate envelope from 20,000 [1/s] to 90,000 [1/s]. These dimensions and rotor speeds covered a range that is common in VADs [12]. Assuming a bulk blood viscosity and density of 3.6 cP and 1.06 g/ml, and taking the fluid gap thickness as the critical length, this corresponds to rotational Reynolds numbers between 115 and 450 in the gap region and Taylor numbers between 170 and 2300. In the axial direction, the blood flow rate was controlled between 5 and 25 ml/min (corresponding to RBC exposure times in the shear gap region between 200 and 1,200 msec and axial Reynolds numbers less than 2).

The axial position of the pump rotor was maintained by passive magnetic forces. The rotor’s radial position was continuously monitored by front and rear magnetic position sensors. These position sensors provide input to a closed-loop feedback controller which modulates the active magnetic bearing system to hold the rotor levitated at the central axis. There is some deviation from the central axis resulting from the magnetic suspension. The front and rear rotor positions are continuously measured via calibrated Hall effect sensors, at 15 kHz through LabView (National Instruments, Austin, TX), and used to extrapolate the radial rotor position in the shearing gap. In this manner, the size of the “orbit” of the rotor is resolved as it deviates from a perfectly centered position, and the size of the shearing gap is measured and used to calculate the magnitude of shear for each blood sample.

Because of the non-Newtonian shear-thinning property of blood, it is important that measurements of blood viscosity be performed at experimentally relevant shear rates. Prior to testing with blood, three Newtonian viscosity standards (1, 3, and 9 cp, AMETEK Brookfield, Middleboro, MA) were passed through the device to calibrate the current drawn by the motor as a function of rotor speed and viscosity (Figure 2). Using this relationship, we were able to use the device to indirectly measure the instantaneous *effective viscosity* via the motor current measurement at different rotational speeds. To avoid undesired thermal effects during calibration, the viscosity standards were pumped through the system at a high flow rate (25 ml/min) corresponding to a maximum of 15 seconds of residency within the device. After calibrating the shearing device with viscosity standards by varying the shearing rate from 24,000 to over 53,000 [1/s] (Figure 2), it was used as an instantaneous high shear viscometer during the blood experiments.

**Figure 2.**
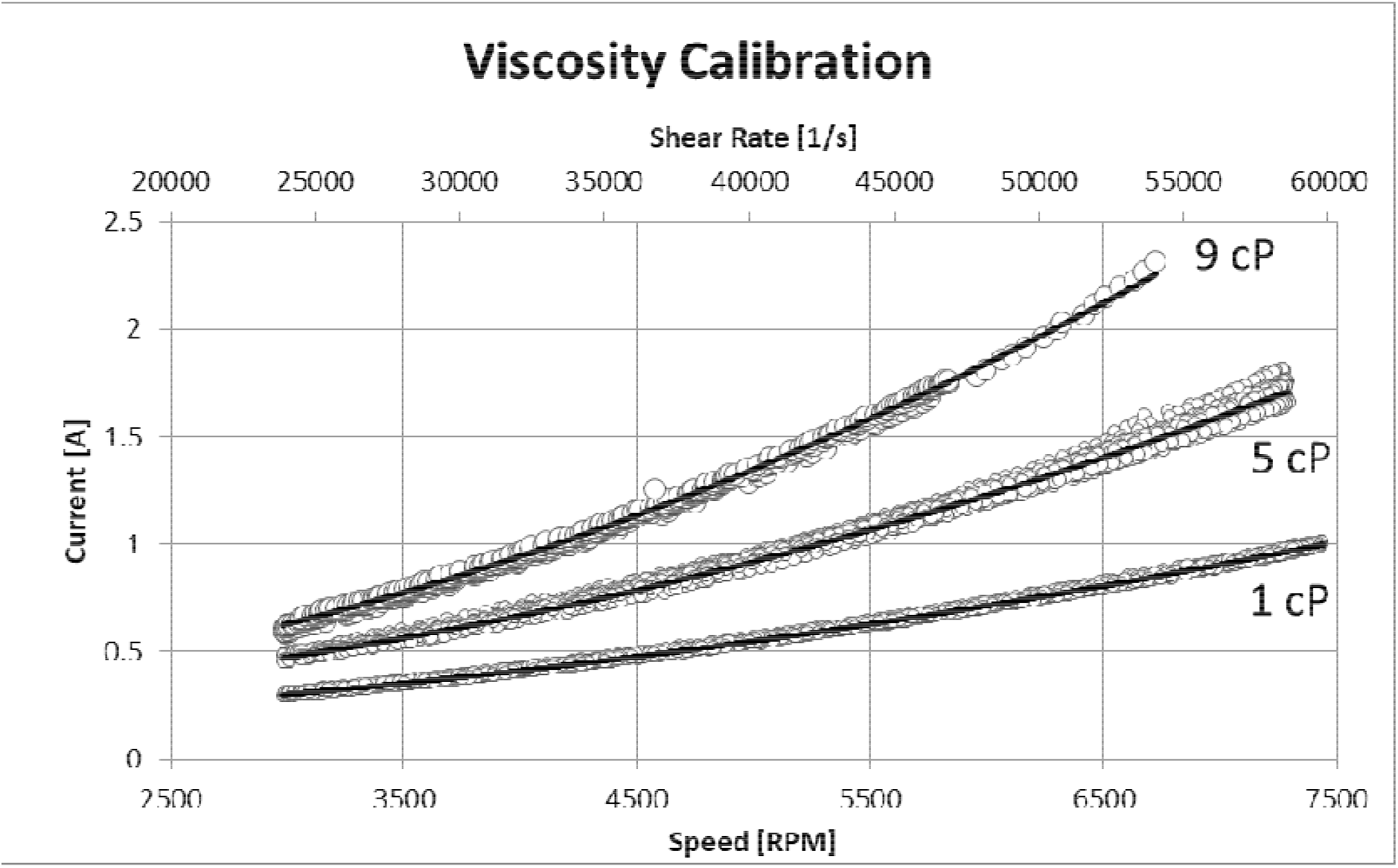
Calibration plot to define the relationship between motor current, rotational speed, shear rate, and viscosity over the same rotational speed sweep used to generate hemolysis curves.

### Test System Characterization

A calibrated syringe pump (PHD Ultra, Harvard Apparatus, Holliston, MA) was used to control flow rate and push blood in a single pass through the shearing device, after which it was collected as a drip from the outlet tubing into 2 ml centrifuge tubes (Figure 3). Pressure was measured at the inlet of the device. Temperature was monitored at the inlet and outlet of the device using K-type wire thermocouples probing into the blood flow. Chilled water (10 °C) was pumped through a custom aluminum cooling “jacket” in order to dissipate the heat generated by the motor and active magnetic bearings.

**Figure 3.**
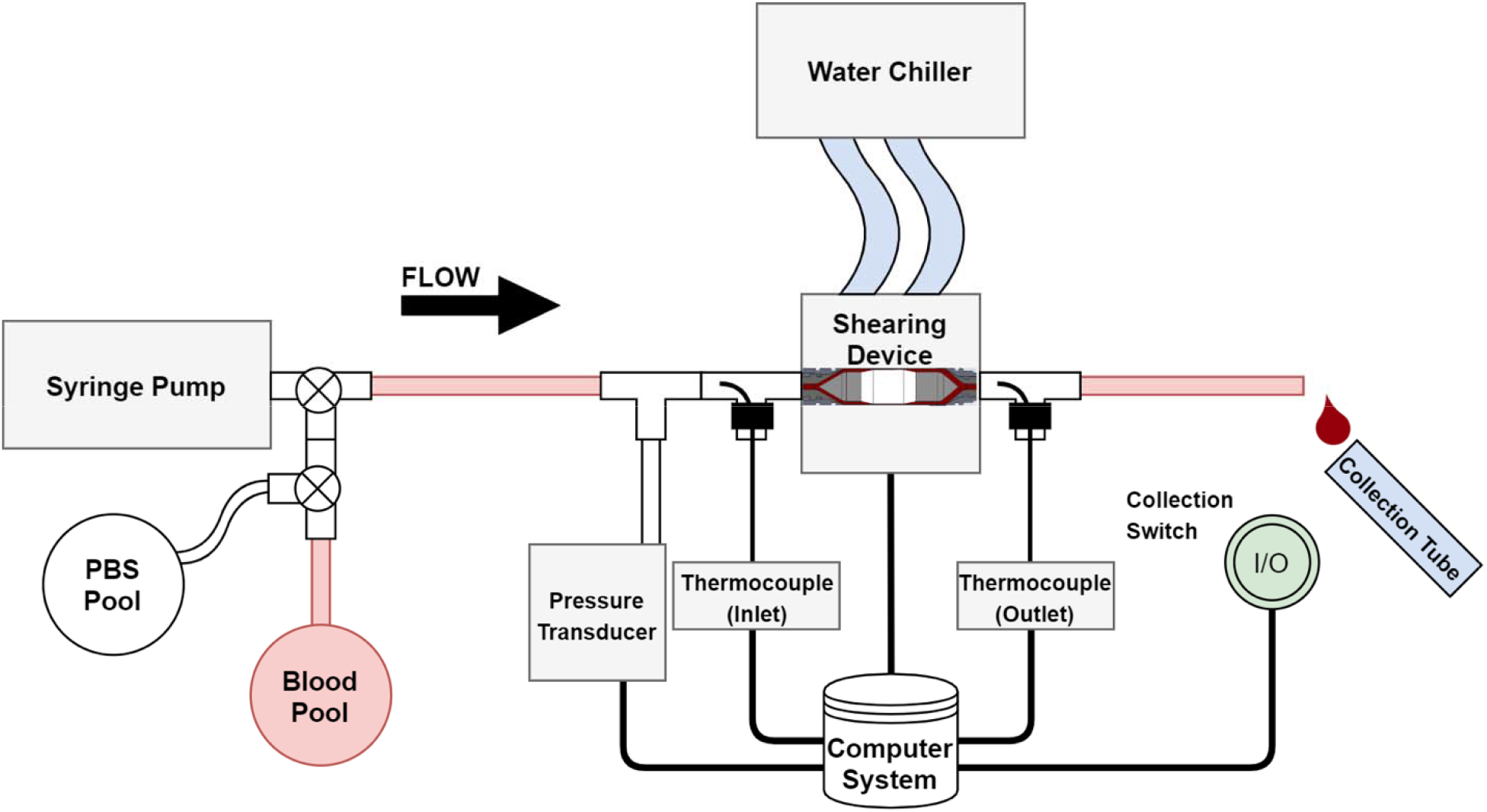
Diagram of experiment apparatus. Shearing device volume is 6 ml. The overall system volume is 8 ml, including inlet and outlet tubing.

At the start of a set of experiments, the system was rinsed with phosphate-buffered saline (PBS) to remove air and impurities and subsequently flushed with gently mixed, room temperature blood. A 60 ml syringe was filled from the same blood pool at a constant withdrawal rate. The syringe pump was set to a constant infusion rate corresponding to a single shearing gap exposure time and the device rotor was idled at its lowest shear rate condition. The entire syringe volume was then infused continuously. After 1.5 times the system fill volume of 8 ml had been purged into a waste container, the outlet blood drip was continuously collected into 2 ml tubes. The rotational speed of the rotor (proportional to the applied shear rate) was ramped from its lowest condition to the device’s maximum RPM over the course of the syringe’s infusion. By repeating this procedure for several different syringe infusion rates, the resulting hemolysis could be mapped over a series of increasing sweeps of shear rate and a range of exposure times with very little wasted blood.

Due to the lag time between when blood was exposed to the shearing gap and when it was collected, a manual data collection switch was used to synchronize and record the system parameters for the individually collected blood samples. At low flow rates, a sample could take several seconds to collect, during which time, the device was slowly increasing its RPM and shear rate. To account for this, as well as the potential blood mixing in the non-gap region of the device, the reported values for each system parameter, including shear rate, are the average values over the time of a 2 ml sample’s residency within the 6 ml test device. The uncertainty in this average is reported as the range of values recorded during this same time interval. This results in a conservative assessment of applied shear for a given sample with some overlap in the uncertainty bounds of consecutive samples, due to the fact that the device volume is 3 times that of a single sample. The uncertainty bounds for a typical sampling interval are the average shear rate ± 5,000 [1/s]. The resulting data are thus to be interpreted in the context of continuous trends or sweeps of increasing shear rate, rather than discrete data.

All data presented from this study had to meet two performance acceptance criteria. Blood samples were discarded if they exited the shearing device with an average temperature outside an acceptable range (23-35 °C), or if the deviation of the rotor orbit exceeded 30 um.

### Blood Handling and Preparation

Porcine whole blood was obtained from a commercial source (Lampire Biological Laboratories, Pipersville, PA) in 1-liter quantities. Donor blood was collected through a 16 Ga needle into a sterile bottle prepared with a 15:85 ratio of ACD anticoagulant to blood and shipped overnight for hemolysis testing less than 24 hours after draw. Upon arrival at the lab, the blood was passed through a 100 μm polypropylene filter mesh. All blood contacting containers were pre-rinsed with PBS. Hematocrit was measured via 75 mm long glass micro-capillary tubes in a hematocrit centrifuge (Microspin 24, Vulcon Technologies, Grandview, MO), and adjusted as necessary via hemodilution with PBS or centrifugal hemoconcentration followed by removal of plasma. Low shear rate bulk viscosity of both whole blood and plasma were measured using a Cannon-Fenske glass tube viscometer (9721-A56, Cannon Instrument Company, State College, PA) at 23 °C and converted from centistokes to centipoise using density, measured gravimetrically with a laboratory balance (XS204, Mettler-Toledo, Columbus, OH). To isolate plasma for the determination of hemoglobin concentration, the collected whole blood samples were centrifuged at 3,000 rcf for 10 min and the resulting supernatant plasma was pipetted into a fresh tube. The plasma was then spun again at 13,000 rcf for 10 min and each sample was aliquoted into three 200 uL replicates in a 96-well plate for absorbance measurement via the Cripps Method to assay free hemoglobin (ΔfHb) as a measure of hemolysis [7]. PBS was used to correct for background absorbance in the spectrophotometric plate reader (Epoch, BioTek Instruments, Winooski, VT) and absorbance values were pathlength corrected to 1 cm. The modified Index of Hemolysis (IH% in Eq. 4) was used to normalize for differences in hematocrit (Hct) and cellular hemoglobin amounts (Total Hb) for each blood experiment [5]. Using the normal range of mean corpuscular hemoglobin concentration for porcine blood as a reference [33], the total blood hemoglobin concentration in g/dL was approximated as 1/3 the hematocrit percentage (e.g. for a hematocrit of 39%, total blood hemoglobin concentration was assumed to be 13 g/dL).

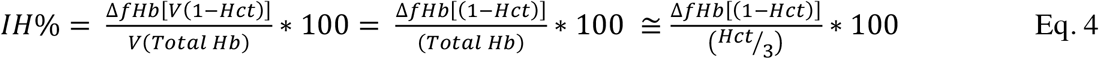

### Experiment 1: Varied Bulk Viscosity via Hematocrit

In order to evaluate the effect of hematocrit on the assessment of shear-induced hemolysis, blood from 3 donors was adjusted into pools with hematocrits ranging from 23% to 41% via dilution with PBS or hemoconcentration. The adjustment of hematocrit served as a means of directly adjusting the whole blood bulk viscosity while maintaining a constant media (plasma) viscosity. Each blood pool was then exposed to a sweep of shear rates from 20,000 [1/s] to 80,000 [1/s] at a blood exposure time of 680 ms (flow rate of 9 ml/min). To compensate for the use of different hematocrit blood pools, the hemolysis levels were normalized using the %IH equation. To evaluate the relationship of shear rate and calculated shear stresses on the hemolysis results, variability in those parameters were assessed at a benchmark hemolysis value of 0.5% IH.

### Experiment 2: Varied Plasma Viscosity via Dextran-40 and PBS

Blood from a single donor was separated into two pools and diluted to hematocrits of 23%, 29%, and 35% using different dilutants. For the first pool, the hemodilution was performed using PBS, and for the second, it was done using a 10% solution of Dextran-40 (31389-25G, Sigma-Aldrich, St. Louis, MO) in PBS. Whereas dilution with PBS maintains media viscosity similar to plasma and decreases bulk viscosity, the added Dextran-40 increased both media viscosity and bulk viscosity. Unlike hematocrit, which changes the bulk viscosity by the addition of more suspended solids, the dextran changes the viscosity of the suspension media directly. Each hematocrit adjusted blood pool was exposed to sweeps of shear rate from 20,000 [1/s] to 80,000 [1/s] at a single blood exposure time of 680 ms. This decoupled the media viscosity from the bulk viscosity and facilitated a direct comparison of the effect of media viscosity on the observed hemolysis at a given shear rate. Where the term “shear stress” is used in Experiment 2, it is calculated as the product of the shear rate and the media viscosity, rather than the more commonly used bulk viscosity, for more direct relevance to the parameters under investigation.

None of the methods involved human subjects or care of vertebrate animals, so no institutional ethical review was required.

## Results

### Experiment 1: Varied Bulk Viscosity via Hematocrit

Over the range of hematocrits evaluated (23% - 40%), the average measured effective bulk blood viscosity ranged from 2 cP to 5 cP, thus a significant variation in applied shear stresses was obtained at the same shear rate conditions. The variability (%CV) in effective viscosity ranged from 6.8% to 11.4% at a given hematocrit and was similar across all donors (Figure 4). The main cause of this variability was attributed to increasing temperature from the active magnetic bearings over the duration of infusion. Per our acceptance criterion, only samples exiting the device with a blood temperature between 23 °C and 35 °C were used in the analysis.

**Figure 4.**
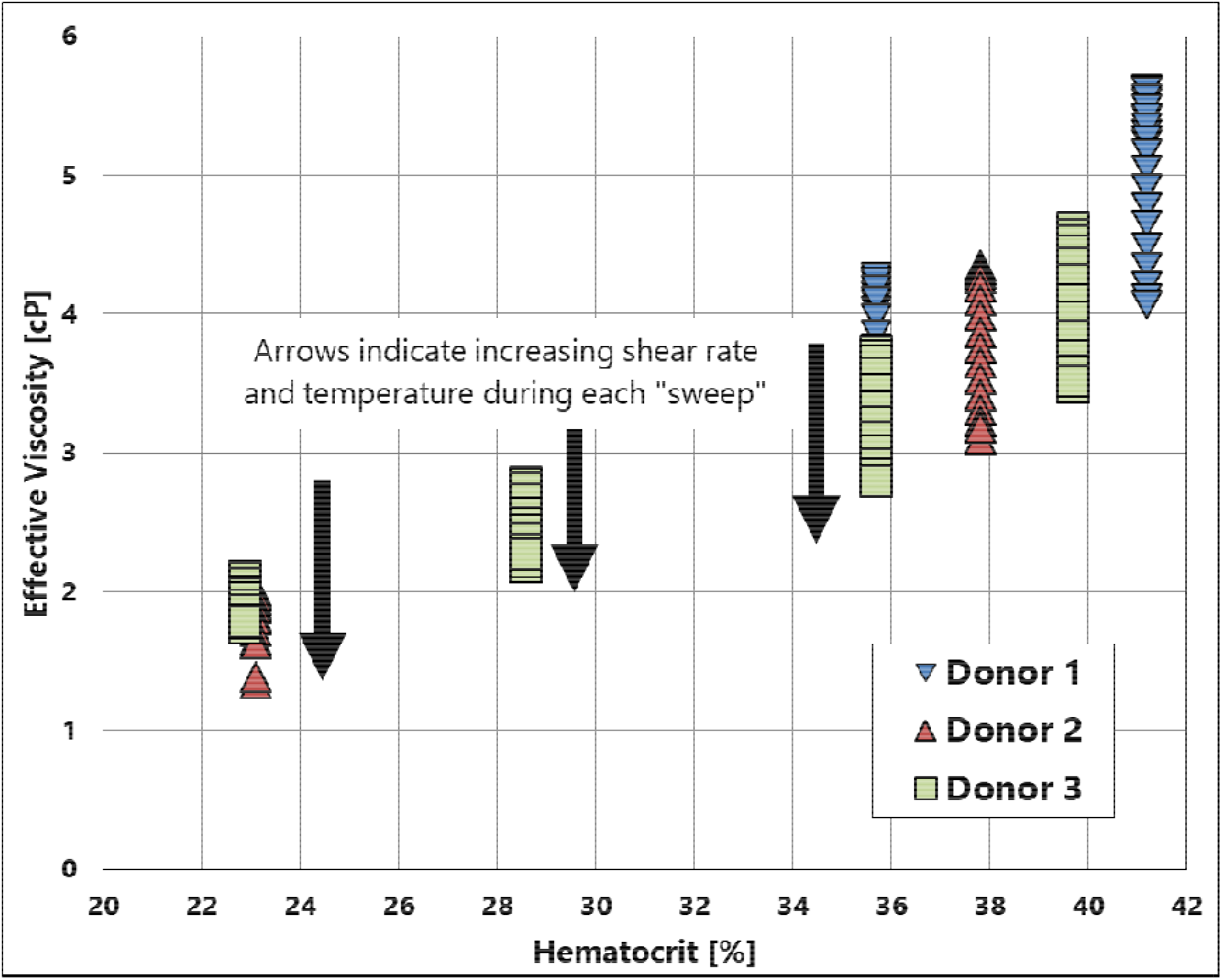
Effective viscosity of porcine whole blood at different hematocrits measured in realtime by the high shear device. Each cluster of data represents a single donor at a given hematocrit over the full range of shear rate used in the experiments. In all cases, the viscosity decreased during the velocity sweep, likely due to the combination of increased shear rate and observed increase in temperature during each run.

After normalizing the plasma hemoglobin concentrations by the different experimental hematocrits, the index of hemolysis for a given shear rate was observed to be similar between each of the seven blood pool tests irrespective of the hematocrit (Figure 5A). Every blood pool reached an IH% benchmark value of 0.5% damage over a narrow range of shear rates (67,500 to 75,000 [1/sec]), which corresponds to an 11% variation). However, when the same data were plotted as a function of shear stress (i.e. the product of shear rate and effective bulk viscosity, Figure 5B), the range of shear stress at the 0.5% IH benchmark value (125 to 300 Pa) was larger (82% of mean) and was dependent on the hematocrit. The variability was similarly high when shear stress was calculated using the low-shear bulk viscosity rather than the high shear effective viscosity. Hence, the resulting hemolysis is better predicted using shear rate alone (Fig. 5A) compared to shear stress as calculated using the bulk viscosity values (Fig. 5B).

**Figure 5.**
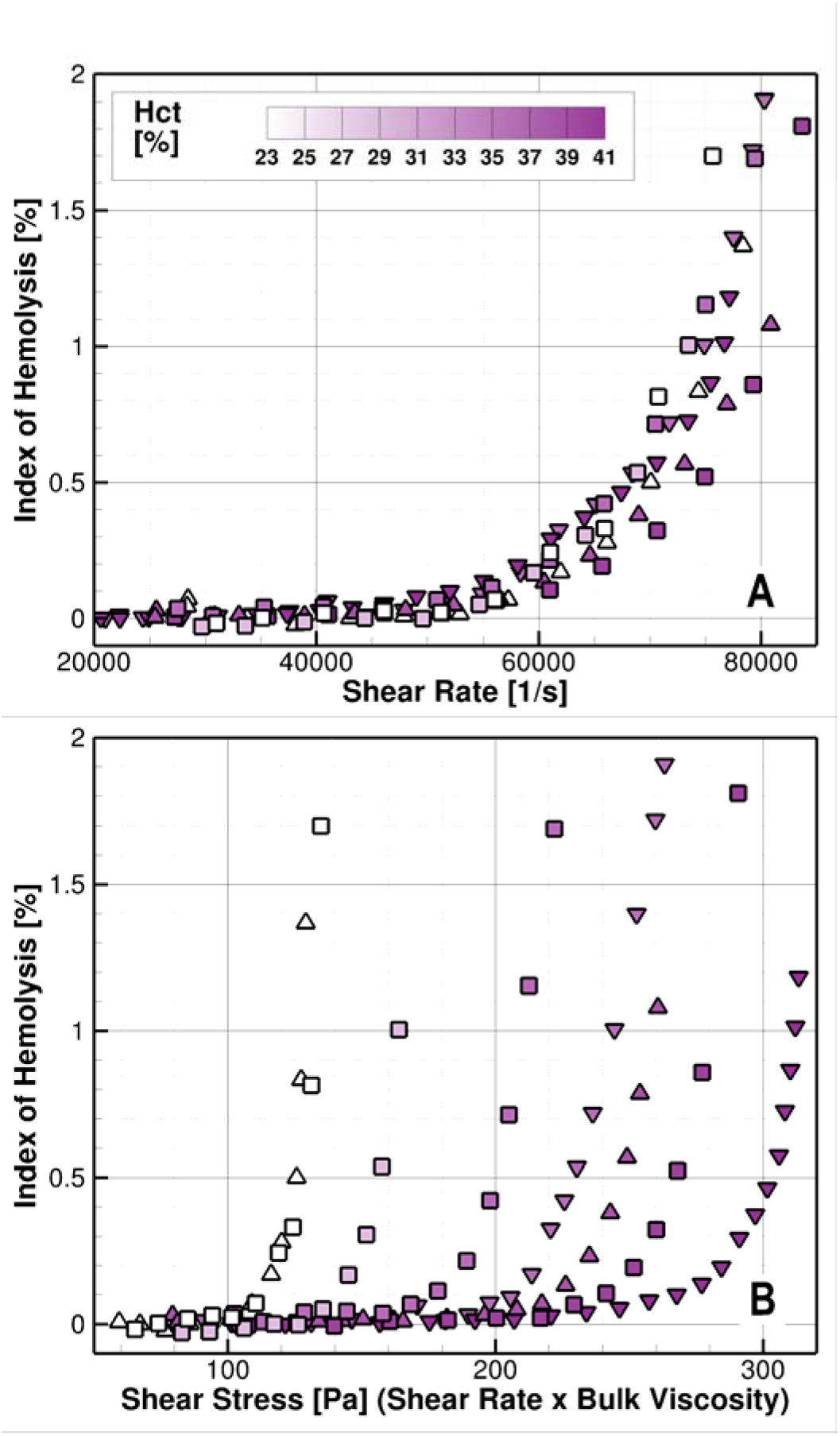
A) Comparison of hemolysis in porcine blood at different hematocrits plotted against the viscosity-independent term shear rate. B) Hemolysis data plotted as a function of the viscosity-dependent term shear stress, as calculated using the bulk fluid viscosity. Blood exposure time was 680 ms.

### Experiment 2: Varied Plasma Viscosity via Dextran-40 and PBS

Hemodilution performed with PBS resulted in proportionally decreased hematocrit and bulk viscosity with little change (from 1.5 cP to 1.2 cP) in media viscosity (Figure 6), as was the case in Experiment 1. Hemodilution with the high viscosity dextran solution, however, resulted in relatively constant bulk viscosity when measured at both low and high shear rates, regardless of hematocrit. Th decrease in viscosity due to the reduction of hematocrit was compensated for by the increase in media viscosity due to the added dextran. For all blood pools, the low-shear rate bulk viscosity measured using the glass capillary viscometer was greater than the effective high-shear rate bulk viscosity measured by the shearing device, consistent with the shear thinning nature of blood.

**Figure 6.**
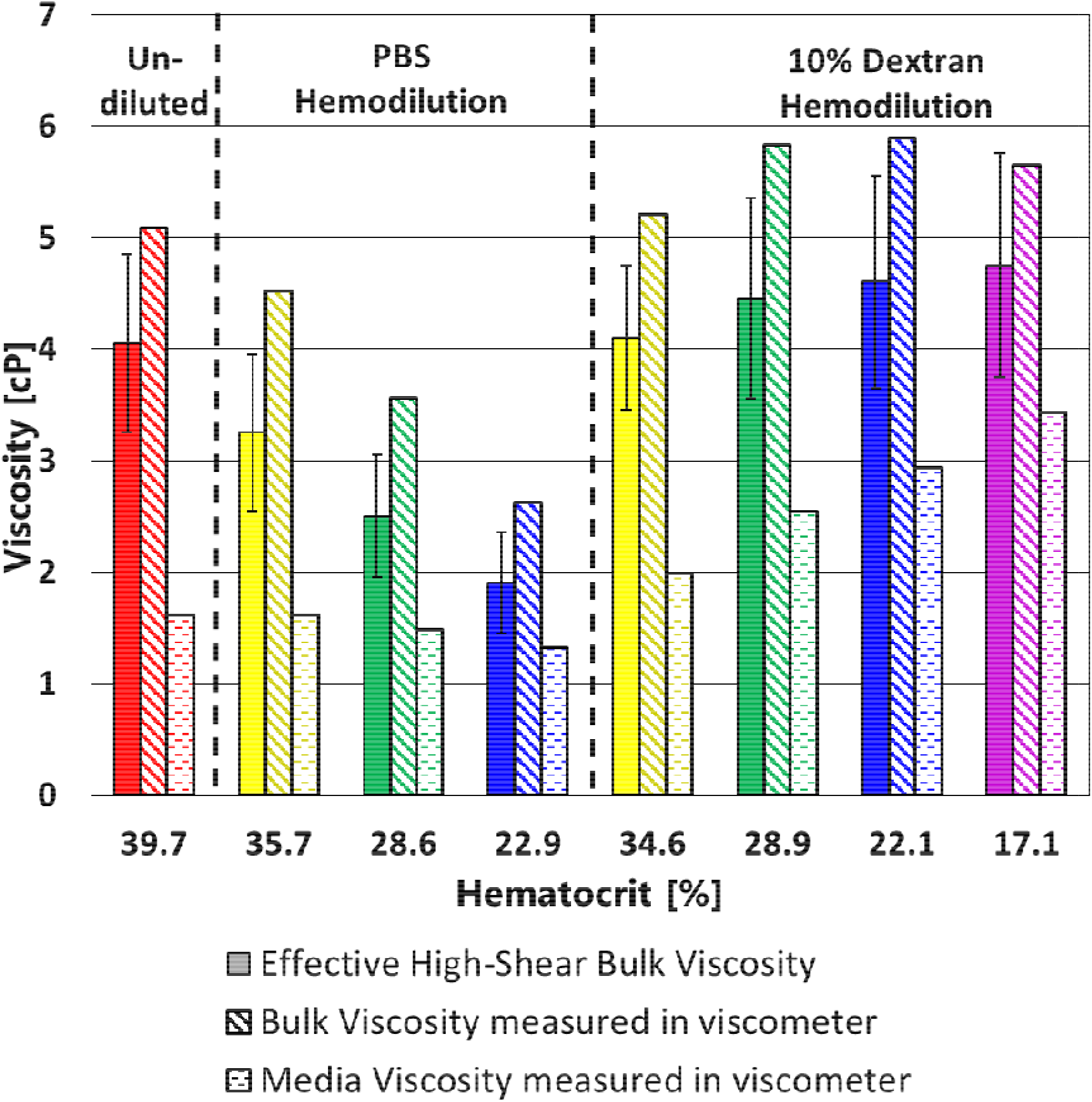
Viscosity of porcine blood components with different dilutants. Color pairs indicate matched hematocrits. Low shear measurements performed using a Cannon-Fenske glass tube viscometer and converted to centipoise. Effective bulk viscosity measured using shearing device.

In Experiment 2, the applied fluid shear stress was calculated as the product of shear rate and media viscosity (rather than bulk blood viscosity) in order to more directly evaluate the parameter being modulated. For this experiment, as was true for Experiment 1, when the index of hemolysis was plotted against shear rate, the data largely followed a single damage curve independent of the hematocrit, media viscosity, or the dilutant used (Figure 7A). The range of shear rates resulting in a 0.5% IH value of damage was again narrow (21% of the mean) and varied between 61,000 and 75,000 [1/sec] for all eight blood pools. Conversely, when the same data were plotted against shear stress (using media viscosity, Figure 7B), a wide (95% of the mean) spread in the data (shear stress ranged from 75 to 210 Pa at the 0.5% IH damage benchmark) was observed (similar to Figure 5B from Experiment 1). Shear stress calculated using effective bulk blood viscosity was similarly variable (Figure 7C), with the exception of an apparent cluster of the dextran hemodiluted blood which all had nearly the same bulk viscosity (Fig. 6). In summation, the resulting hemolysis values in Experiment 2 were more strongly correlated to shear rate (Fig. 7A) than to shear stress (Figs. 7B or 7C) as calculated using either the media or the effective bulk whole blood viscosity.

**Figure 7.**
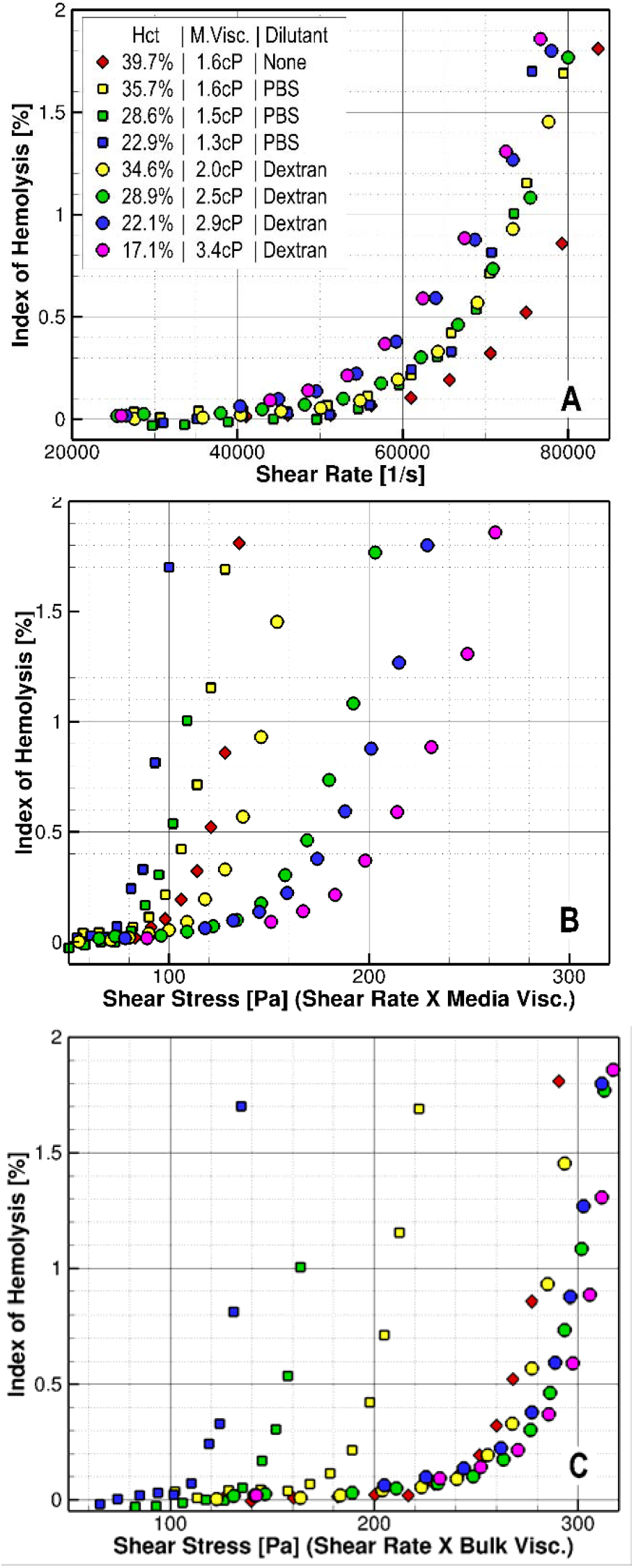
A) Comparison of hemolysis in porcine blood of different media viscosities plotted against the viscosity-independent term shear rate. B) Hemolysis data plotted against the viscosity-dependent term shear stress, here calculated using the media viscosity. C) Hemolysis data plotted against the viscosity-dependent term shear stress, here calculated using the effective high-shear bulk viscosity. Blood exposure time was 680 ms. Color pairs indicate matched hematocrits.

## Discussion

As summarized by Leverett et al. [9], fluid shear stresses and RBC exposure times have been recognized as key parameters in blood damage modeling experiments for nearly 50 years. The power-law formulation derived by Giersiepen et al. is foundational and widely referenced in fitting empirical hemolysis data from experiments on different test devices [8] [34] [24] [35]. It is also the most common equation form for numerical models to assess blood damage used in design processes [36][37][38]. The power-law model asserts that hemolysis is proportional to the product of exponential functions of both fluid shear stress and exposure time, as given below where C, α, and β are constants derived from a fit to experimentally acquired data.

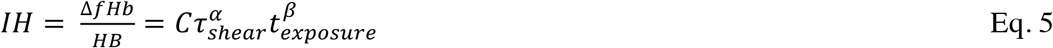

For both experimental and computational models, the shear stress in this equation is usually calculated by multiplying the shear strain rate by the bulk viscosity of whole blood [21]. Although blood is non-Newtonian, in some cases the bulk viscosity is measured at extremely low shear rates and not representative of high shear flow regions where researchers are most concerned with mechanical RBC damage [13]. In other cases, the bulk viscosity is not measured at all, but rather approximated as a constant based on published values or relationships between viscosity and hematocrit. Recently, Goubergrits, et al. noted that difficulty in accurately measuring blood rheological properties over a wide range of shearing rates is one detriment to developing computational predictive models of blood damage in medical devices [39]. A unique feature of our shearing device is the ability to estimate the effective bulk viscosity at each test condition over a shear rate range of 20,000 to 90,000 [1/s]. This allowed us to examine the effects of both shear stress and shear rate over a realistic range experienced by RBCs in clinical VADs, and to address whether bulk blood viscosity or media viscosity had a greater impact on predicting hemolysis.

Experiment 1 challenges the understanding that RBC damage is best represented as a function of shear stress, rather than shear rate. Hemolysis data presented in Figure 5A show that, even across a wide range of physiological hematocrits and corresponding range of bulk viscosities in our Couette flow system, RBC damage (normalized by taking into account the different hematocrits and total hemoglobin concentrations in each blood sample) was highly correlated with the shear rate. The conclusion that damage is more a function of shear rate than shear stress is further reinforced by the analysis of Figure 5B. When the hemolysis level was plotted as a function of shear stress (calculated as the product of shear rate and its corresponding effective bulk viscosity) there was not a clear relationship. If Giersiepen’s formulation of damage as dependent on shear stress was accurate for a range of hematocrits, one would expect the hemolysis data presented in Figure 5B to collapse to a single curve, whereas the shear rate data in Figure 5A would show significant variation. However, the data demonstrate an opposite effect. This lack of dependence of damage on the bulk viscosity (here modulated by hematocrit) suggests that the damage is more a function of shear rate rather than shear stress. By extension, the traditional method of calculating shear stress as the product of bulk viscosity and shear rate further complicates the analysis as it introduces additional uncertainty related to determining an accurate local viscosity. For these reasons, it would be simpler, and probably more accurate, to evaluate blood damage based on the shear rate rather than the shear stress.

Using shear rate instead of shear stress seems reasonable as mechanical hemolysis should occur as a result of local strain on the cell membrane which results from the cell-media interactions. Analytical [40] and numerical [41] models of RBC tank treading predict shear stresses on the surface of the cell between 2.5 and 4 times the product of plasma viscosity and bulk shear rate [28]. This proportionality to shear rate with no consideration of bulk viscosity is consistent with our findings in experiment 1. The effect of cell-cell interactions, which would lead to increased damage at higher hematocrits, was not evident in our study. This is consistent with Brooks et al. [27], who indicated that for a laminar high shear flow, cell-medium stresses dominate cell-cell interactions. Thus, the shear stress calculated by using the bulk viscosity would not capture the hemolytic response of RBCs as effectively as the local fluid shear rate would.

While Experiment 1 investigated the effect of varying hematocrit and bulk viscosity, Experiment 2 was conducted to evaluate the effect of varying the viscosity of the suspension media (plasma with and without Dextran-40) on the resultant hemolysis. As was the case for Experiment 1, the results suggest that shear rate alone, with no consideration of the viscosity of the fluid, correlates with the resulting RBC damage, even across a wide range of media viscosities (1.3 to 3.4 cP), as shown in Figure 7A. Morris and Williams [42] found a minimal effect of suspension media viscosity (also varied by Dextran) below 5 cP on RBC deformability and hemolysis, but a strong affect at higher (6 - 20 cP) media viscosities. Unlike our experiments, Morris and Williams’ high suspension media viscosities resulted in supraphysiological bulk viscosities. Our attempt to model hemolysis as the result of local shear stress, as calculated by the shear rate and media viscosity (from 1.2 to 3.3 cP), was significantly worse than just using the shear rate and neglecting viscosity (i.e. hemolysis in Figures 7B and 7C show similar variation with shear stress as in Figure 5B). This suggests that the formulation of cell membrane stress as the product of local shear rate and the media viscosity is also not valid within the media viscosity range examined here. Experiment 2 further supports that hemolysis was primarily a function of shear rate in our Couette flow model, rather than a viscosity-dependent shear stress, no matter whether local media or bulk viscosity was used to calculate shear stress.

Using shear rate instead of shear stress can also be supported mechanistically. The simplest mechanical analyses of a fluid stress applying a load to a cell assumes that the cell behaves as a static object with a constant orientation, in which case membrane stress may be proportional to fluid stress. However, RBCs are highly deformable and can take multiple shapes [13]. They can align in the direction of flow, tumble and collide with each other, and exhibit “tank-treading” motion in which the membrane rotates over their intracellular contents [13] [43]-[48]. Any or all of these characteristics could potentially absorb energy and dissipate the load that the suspension media would apply to a cell. In the extreme scenario of a spherical particle spinning within a shear flow, there may be minimal membrane stress as there is very little velocity difference between the membrane and surrounding fluid. With all this in mind, it seems likely that there may be some dependence of hemolysis upon the viscosity of the suspension media, but this may be a second-order effect such that membrane stress and the resulting damage are not linearly proportional to fluid stress. Figure 7A hints at this, suggesting that higher viscosity suspension media results in slightly more damage, when compared at a fixed shear rate. Interestingly, this is the opposite conclusion one might draw if considering damage as a function of fluid shear stress using the media viscosity (Figures 7B and 7C), namely that cells can withstand higher stress if the media viscosity is higher. We reject this conclusion and believe that this emphasizes the importance of our two experiments that effectively decoupled shear rate from calculations of shear stress and the effects of media viscosity and hematocrit on bulk whole blood viscosity.

A key piece of the hemolysis puzzle for modeling and predicting blood damage is to determine how fluid stress translates into actual RBC membrane strain [1,13,14,43]. This is a challenge as RBCs continually deform, elongate, tumble, and tank-tread in different flow regimes; therefore, it is important to experimentally investigate the fluid parameters and damage models (e.g. power-law, energy dissipation rate) which may better predict hemolysis. In a recent comprehensive review of experimental and computational models of flow-induced hemolysis, Faghih and Sharp point out that it is still unclear whether bulk fluid or local media viscosity should be used to represent the transfer of fluid stresses to the RBC membrane [14]. The results for our experimental model suggest that shear rate is the dominant factor leading to hemolysis, independent of the viscosity used. Other researchers have already developed experimental models and reported results based on blood shear rate rather than shear stress [49].

The consequences of our findings may be significant. Although some studies present results in terms of shear rate [50], most blood damage studies calculate the applied RBC stress as the product of a measured or theoretical shear rate and a single measurement or reference value for whole blood viscosity, and then compare it to published shear stress thresholds for hemolysis. The conclusions of these prior studies are therefore dependent on the chosen value for whole blood viscosity. In order to maintain a constant bulk fluid shear stress, other researchers have adjusted the experimentally applied shear rate in order to compensate for variation in bulk viscosity (due to hematocrit or temperature, etc.) [13]. As such, in an effort to report results at an equivalent shear stress, they may have inadvertently confounded the experimental conditions by forcing a mismatch of shear rate. The findings of the current work, particularly of Experiment 1, suggest that the common approach of calculating stress from shear rate could complicate the observed relationship of applied shear stress to hemolysis and potentially influence study conclusions. This is especially problematic as the values for the viscosity, shear rate, and shear stress for each individual experiment are rarely reported.

Although our findings were largely consistent for our test device, we present four limitations of this experiment that should be considered when interpreting or generalizing conclusions drawn from these results. First, it is likely that the non-gap region of the device under a Taylor-Couette flow regime contains Taylor vortices, which could contribute to some mixing between sequentially collected blood samples. However, this is mitigated by reporting each test parameter as the average value recorded during a 2 ml sample’s duration of residency within the 6 ml device, as described in the methods. While this increases the uncertainty in fluid shear experienced by a red blood cell, it does so in a well characterized and repeatable manner due to the highly automated protocol. Even if this were to affect absolute values of IH%, it would have no effect on the trends between blood pools, nor on the resulting conclusions comparing shear stress to strain rate. Similar to the flow regime investigated in our device, the possibility of Taylor vortices may also be present in rotary flow VAD devices.

Second, although the magnetic levitation system reduces the complexity of the flow path, and removes high shear regions around conventional mechanical bearings, it does introduce other considerations – i.e. radial deviation of the rotor due to the magnetic levitation as well as heat contributed by the magnetic bearing system. As described above, these tertiary factors were directly measured and data were excluded from the study when the blood temperature or rotor orbit deviation exceeded allowable limits. These exclusions only occurred when the device was operating at extremely high shear rates; experimental conditions below 70,000 [1/s] were unaffected and all of the conclusions made in this work remain supported even if we limited our observations to a lower range of shear rates.

A third limitation was that once gently mixed blood from the pool was loaded into the injection syringe and placed in the pump, the syringe was kept in a horizontal position for the duration of infusion through the shearing device – approximately 7 min. During this time, it is possible that RBCs began to settle and separate from the plasma in the syringe, although no such separation was observed by visual inspection during the experiments. If cell separation were to occur in this apparatus, it would be mitigated by the horizontal configuration of the infusion syringe and would likely only become a notable factor towards the end of the duration of infusion. Moreover, all blood pools would be affected similarly, as they had the same blood volume and infusion rate. Cell separation, however, is always an important consideration when conducting testing with whole blood, particularly when studying the effects of hematocrit.

We are not the first to use Dextran for the purpose of modifying media and bulk viscosities in order to study RBC mechanics [16], [27], [42], [43], [45], [48], [51]. A fourth limitation was that our study, and those prior, assume that the addition of Dextran to modify the plasma viscosity does not affect the mechanical properties and fragility of RBCs and their cell membranes. Additionally, it is important to note that there is a high degree of donor to donor variability in blood fragility that has been reported in the literature [2]–[4]. However, despite investigating a small number of porcine donors, the conclusions from Experiment 1 (n=3 donors) and Experiment 2 (n=1) are unlikely to change from donor to donor. This is because the phenomena studied pertains to the mechanism by which fluid shear imparts RBC membrane strain in relationship to bulk blood viscosity and local media viscosity. While RBC fragility may be variable between donors and species [13], the fundamental mechanism of RBC damage under investigation is likely universal. However, a more expansive parametric study using human and animal blood is suggested to explore these findings.

Finally, further study of the relationships observed by this work would be well-served by more direct analysis of the mechanistic relationship between shear and extensional flows, and the effect of exposure time, to the deformation of RBCs when examined in whole blood. This work could also be expanded to investigate different flow regimes besides the laminar Taylor-Couette flow field studied here (e.g. transitional, pulsatile, and turbulent flows).

In conclusion, we found that the fluid shear rate in our experiments captured the hemolytic characteristics of blood better than either bulk or media viscosity-based formulations of shear stress. These findings may have considerable ramifications for the commonly used approaches, such as the power-law model, to estimate hemolysis from results of computational simulations or empirical measurements. This is especially true for experiments where the applied shear rate has been adjusted to maintain a constant applied shear stress by accounting for changes in bulk blood viscosity. As this could compromise the results, it is recommended that future investigations of shear-induced blood damage report findings with respect to the viscosity-neutral term of shear rate, in addition to reporting the bulk whole blood viscosity measured at an appropriately high shear rate, and plasma viscosity.

## Acknowledgements

We thank Dr. Luke Herbertson of the Center for Devices and Radiological Health at the FDA for reviewing the manuscript. This project was supported in part by a student appointment [JK] to the Research Participation Program at the FDA, administered by the Oak Ridge Institute for Science and Education through an interagency agreement between the U.S. Department of Energy and the FDA, with additional student support from the Harvey J Palmer Endowment to RIT’s Department of Biomedical Engineering.

## (iv)

There are no tables in this manuscript

